# The effect of metal remediation on the virulence and antimicrobial resistance of the opportunistic pathogen *Pseudomonas aeruginosa*

**DOI:** 10.1101/2022.09.20.508257

**Authors:** Luke Lear, Elze Hesse, Laura Newsome, William Gaze, Angus Buckling, Michiel Vos

## Abstract

Metal contamination poses both a direct threat to human health as well as an indirect threat through its potential to affect bacterial pathogens. Metals can not only co-select for antibiotic resistance, but also might affect pathogen virulence via increased siderophore production. Siderophores are extracellular compounds released to increase ferric iron uptake — a common limiting factor for pathogen growth within hosts – making them an important virulence factor. However, siderophores can also be positively selected for to detoxify non-ferrous metals, and consequently metal stress can potentially increase bacterial virulence. Anthropogenic methods to remediate environmental metal contamination commonly involve amendment with lime-containing materials, but whether this reduces *in situ* co-selection for antibiotic resistance and virulence remains unknown. Here, using microcosms containing metal-contaminated river water and sediment, we experimentally test whether metal remediation by liming reduces co-selection for these traits in the opportunistic pathogen *Pseudomonas aeruginosa* embedded within a natural microbial community. To test for the effects of environmental structure, which can impact siderophore production, microcosms were incubated under either static or shaking conditions. Evolved *P. aeruginosa* populations had greater fitness in the presence of toxic concentrations of copper than the ancestral strain, but this effect was reduced in the limed treatments. Evolved *P. aeruginosa* populations showed increased resistance to the clinically-relevant antibiotics apramycin, cefotaxime, and trimethoprim, regardless of lime addition or environmental structure. Although we found virulence to be significantly associated with siderophore production, neither virulence nor siderophore production significantly differed between the four treatments. We therefore demonstrate that although remediation via liming reduced the strength of selection for metal resistance mechanisms, it did not mitigate metal-imposed selection for antibiotic resistance or virulence in *P. aeruginosa*. Consequently, metal-contaminated environments may select for antibiotic resistance and virulence traits even when treated with lime.

**Graphical abstract:** 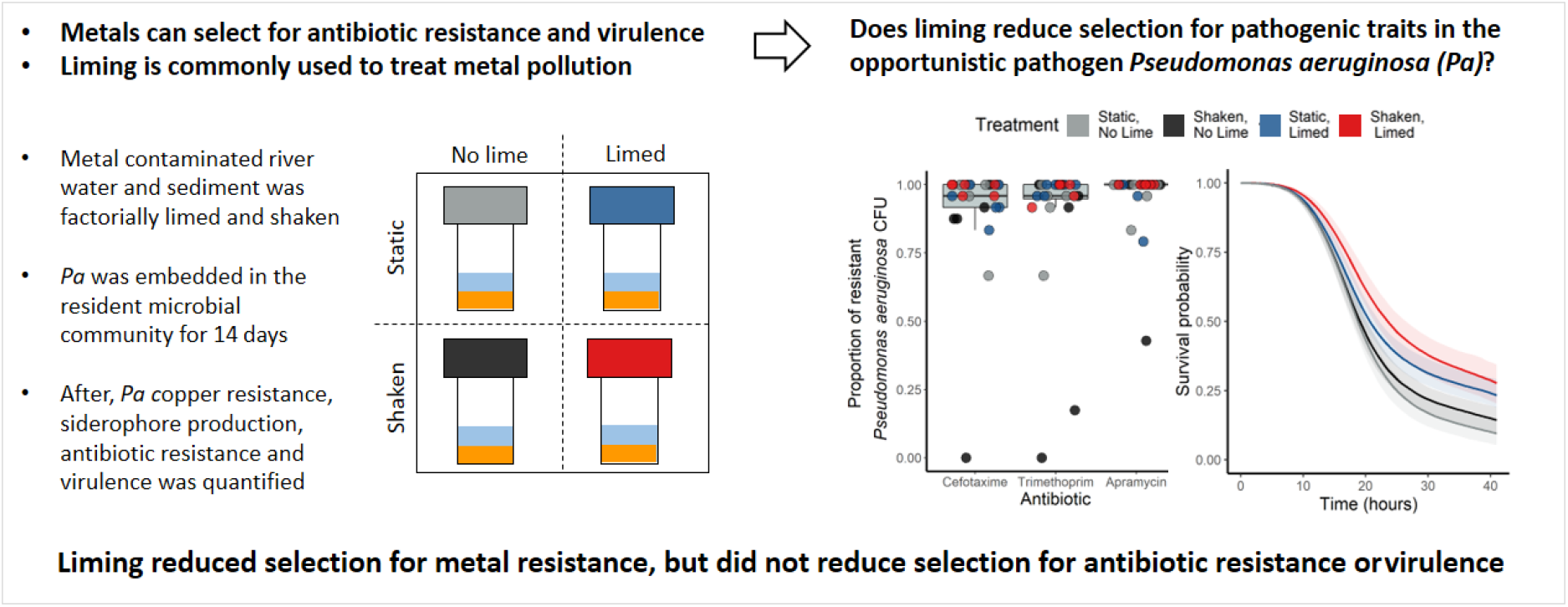

## Introduction

Metals are ubiquitous in the Earth’s crust and many are essential for cellular processes (1-3). However, agriculture and industry have resulted in toxic levels of metal contamination in many environments via pesticide use, sewage sludge application, atmospheric deposition and mining (1, 4). Human exposure to metals via contaminated crops or water poses a serious health threat (5-9). Furthermore, metal contamination can decrease microbial diversity, biomass and functionality, which in turn can affect ecosystem function (10-12). Such are the potential health and economic costs that metal remediation is common practice (13). Often, this involves adding lime-containing materials to acidic metal-contaminated environments (14-16). By raising the pH, liming causes metal ions to precipitate and become less soluble and consequently less bioavailable (10, 13, 17-19). This decreases metal uptake by plants which helps alleviate phytotoxicity and prevent metals entering the food chain (20). Lime addition is the oldest and most widely used metal remediation method (19), and has been used to treat contamination in lakes (14, 21), rivers (22) and soils (15, 16, 18).

Reducing metal bioavailability by liming will inevitably affect the microbial community (18, 23, 24). However, how such remediation could affect the ability of environmental pathogens to cause infection or to withstand antibiotic treatment remains largely unexplored. A key microbial trait likely to change after liming is the production of siderophore compounds (18). The canonical function of siderophores is to aid iron (Fe) sequestration from the extra-cellular environment (25, 26). Fe is vital for microbial growth as a cofactor for a number of essential enzymes (27, 28), but is most commonly present as insoluble Fe^3+^ and therefore is of limited bioavailability, especially at near-neutral pH (27-32). Siderophores are released by cells where they form extracellular complexes with Fe^3+^, these are then taken up by selective outer-membrane transport proteins before Fe^3+^ is reduced to bioavailable Fe^2+^ and the siderophore made available for reuse (33). Siderophores are important virulence factors as they allow pathogens to grow within hosts that actively withhold iron (34, 35). Apart from iron, siderophores can also chelate toxic metal ions, but these complexes cannot re-enter the cell due to the selectivity of the outer-membrane transport proteins (28, 29). This means siderophore production can be selected for as a detoxifying method in the presence of bioavailable toxic metals (26, 29, 36). Consequently, toxic metal concentrations can select for greater virulence by selecting for increased siderophore production (37). Lime remediation of metal-contaminated environments thus could potentially select either for the upregulation of siderophore production when it predominantly results in decreased bioavailability of Fe, or for the downregulation of siderophore production when it predominantly results in lower metal toxicity, with concomitant expected changes in virulence. Previous work has shown siderophore production to decrease as a consequence of liming at the level of whole microbial communities (18), but whether this also occurs in environmental pathogens that rely on siderophore-mediated iron uptake remains untested.

It is well established that some mechanisms that bacteria use to resist metal contamination also confer resistance to antibiotics (38). This can occur through cross-resistance when a single mechanism provides resistance to both types of stressors (e.g. efflux pumps (38-44)), through co-resistance when metal and antibiotic resistance genes are located on the same genetic element (45, 46)), or through co-regulation when transcriptional and translational responses to both stressors are linked (38, 43, 47-49). However, to our knowledge, it remains untested whether metal remediation could decrease such co-selection for antibiotic resistance.

In this study, we use the opportunistic pathogen *Pseudomonas aeruginosa* to test whether liming alters virulence by influencing siderophore production, and whether it decreases co-selection by metals for antibiotic resistance. We applied an experimental evolution approach, utilising microcosms containing water and sediment and the resident microbial community from a river heavily contaminated with historical mine waste (50, 51). We embedded *P. aeruginosa* within this natural microbial community and quantified antibiotic resistance, siderophore production and virulence in this focal species after 14 days. *P. aeruginosa* is responsible for a significant proportion of nosocomial infections, particularly those in intensive care units and immunocompromised patients (52). This species is of significant clinical importance as it is resistant to many treatments, both intrinsically and due to its ability to rapidly evolve resistance (53). Outside of the clinical setting, *P. aeruginosa* is commonly found in soil and water (54). The production of siderophores by *P. aeruginosa* is well-studied as a virulence factor, metal resistance mechanism and public good (25, 27, 29, 55-57). Furthermore, the growing interest in its use along with other siderophore producing species to assist phytoremediation of metals using plants (28, 58), makes it an ideal focal species for this study.

The insect infection model *Galleria mellonella* (Greater Wax Moth larvae), a low-cost and ethically expedient alternative for mammalian virulence screens (59), is used here to quantify *P. aeruginosa* virulence (60). We quantified total siderophore production using a CAS assay (61) and pyoverdine production – the main siderophore produced by *P. aeruginosa* (62) – by measuring fluorescence; and tested whether these are correlated with virulence. Extracellular siderophore-metal complexes offer a fitness advantage not only to the producer but also to neighbouring cells, whether these are fellow-producers or not (63-65). Non-siderophore producing ‘cheats’ could gain a selective advantage as they benefit from siderophore production but do not carry the cost of production (25, 30, 31, 66). Cheat fitness is increased in spatially unstructured environments because the greater mixing increases the opportunity to take up siderophore-iron complexes and benefit from siderophores detoxifying the area (65). To take into account the effect of spatial structure on siderophore production, and consequently virulence, we performed our experiments in both static and shaken microcosms. We tested whether the addition of lime or a change in spatial structure affects *P. aeruginosa* resistance to the antibiotics apramycin, cefotaxime and trimethoprim. Both apramycin and cefotaxime have been declared ‘critically important’ for human medicine, and trimethoprim ‘highly important’ by the WHO (67). Moreover, apramycin has been shown to be effective against highly drug resistant strains of *P. aeruginosa* (68).

## Methods

### Collection of river samples and microcosm set-up

Water and sediment samples were collected from the metal-contaminated Carnon River in Cornwall, U.K. (50°13’54.6’ N, 5°07’48.7’ W) (53). We chose this site as it is polluted by multiple toxic metals (18, 50) and liming has been shown to reduce total non-ferrous metal availability in samples collected in the near vicinity (18). The water sample in reference (50) was taken at the same time as the samples used in this study, and contains high concentrations of non-ferrous metals. Sediment was collected using a sterile spatula and water was collected by filling a sterile 1000 mL duran bottle (Schott Duran, Munich, Germany). Sediment (3 g +/-0.1 g) and river water (6 mL) was added to each microcosm (25 mL, Kartell, Noviglio, Italy). The combined water and sediment pH was measured using a Jenway 3510 pH meter (Jenway, Essex, UK).

### Experimental design

Two treatments – liming (lime amendment/no amendment) and spatial structure (shaken/unshaken) – were carried out in a full factorial design (Fig. 1); six replicates were used per unique treatment combination resulting in a total of 24 microcosms. All microcosms were incubated at 20°C. To raise the pH from 5.8 to ∼7.0 to represent a metal remediation scenario, 30 mg (+/- 1.0 mg) of undissolved hydrated lime (Verve Garden lime, Eastleigh, U.K. (18)) was added to each relevant microcosm, then left for 14 days to equilibrate. To observe differences between structured and non-structured environments, microcosms were either kept static or were continuously shaken at 210 rpm (Stuart orbital incubator S1600, Staffordshire, UK). Shaking began on day 14 and ended on day 28 (Fig.1).

**Figure 1.**
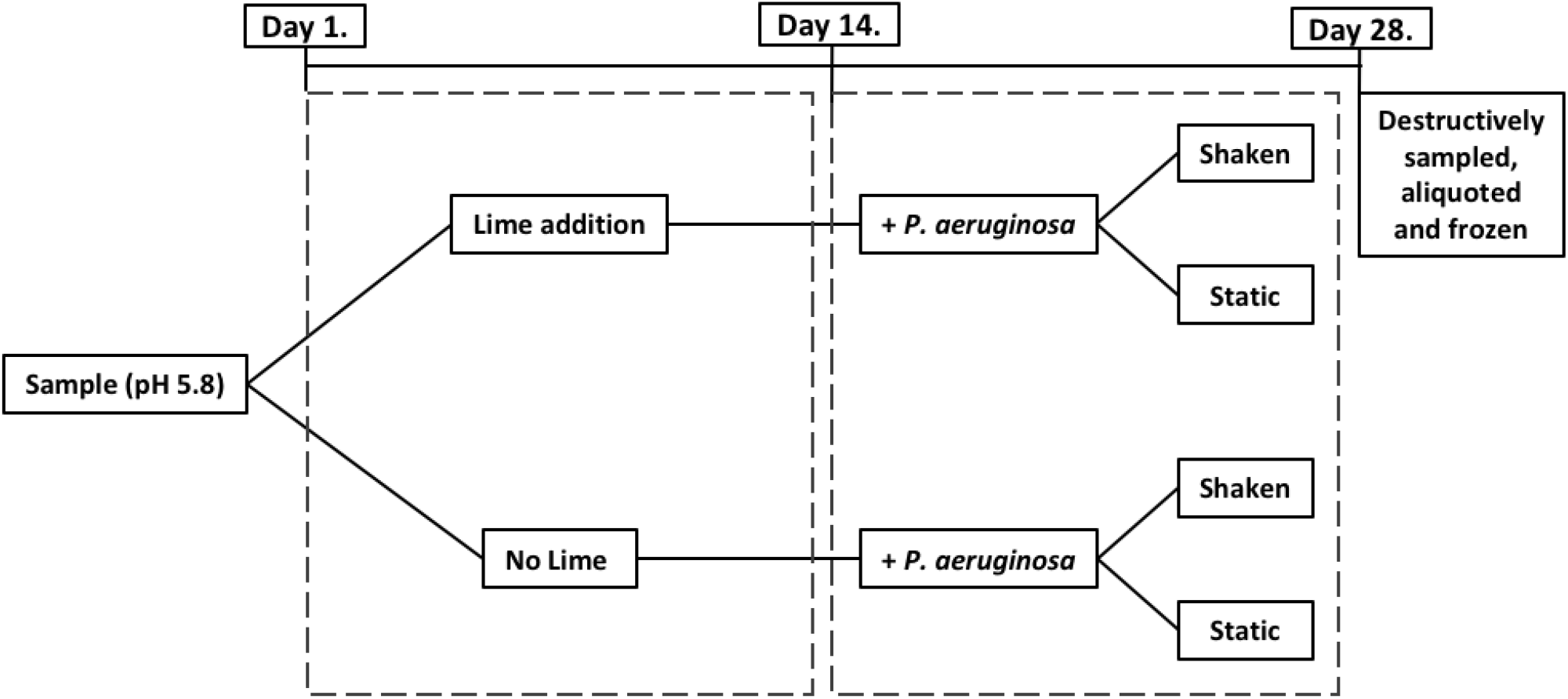
Timeline of experimental design. Lime addition and shaking treatments were performed factorially, with lime added on day 1 and shaking at 210 rpm starting on day 14. *Pseudomonas aeruginosa* was added to all microcosms on day 14. Microcosms were destructively sampled on day 28. Six replicates were used for all treatments (24 microcosms in total).

On day 14, 30 μL (7.3 × 10^8^ colony forming units: cfu) of *Pseudomonas aeruginosa* (PAO1^R^*lacZ*: (69)) was added to each microcosm. This lab-strain is both lacZ marked and gentamicin resistant allowing it to be easily distinguished from the rest of the community on agar containing X-gal (5-bromo-4-chloro-3-indolyl-β-D-galactopyranoside; 100 μg/L; VWR Chemicals) and gentamicin (30 μg/mL Sigma). *P. aeruginosa* was grown overnight in shaking microcosms containing 6 mL of King’s medium B (KB; 10g glycerol, 20g proteose peptone no. 3, 1.5 g K_2_HPO_4_,1.5 g MgSO_4_, per litre). To remove residual nutrients, cultures were centrifuged at 3500 rpm (1233 g) for 30 minutes, after which supernatant was decanted, and the pellet resuspended in half the volume of M9 salt buffer (3 g KH_2_PO_4_, 6 g Na_2_HPO_4_, 5 g NaCl/L) followed by plating on KB agar to calculate the inoculation density. On day 28, all microcosms were destructively sampled by adding sterile glass beads and 12 mL of M9 buffer and vortexing for one minute. Samples were then aliquoted and stored in glycerol (25% final volume) at -80°C.

### Iron analysis (ferrozine assay)

To determine if liming affected Fe speciation and therefore bioavailability, a ferrozine assay was used to measure relative concentrations of Fe^2+^ and total bioavailable iron (70, 71). The first step of this assay quantifies Fe^2+^ which is easily obtainable by bacteria and so does not require siderophores. The second step quantifies both Fe^2+^ and Fe^3+^ and therefore gives a measure of total bioavailable iron including that which requires scavenging mechanisms, such as siderophores. By dividing the first measurement by the second it is possible to estimate the proportion of iron in each treatment that is of relatively high bioavailability to *P. aeruginosa* (70, 71). The first measurement is given by digesting 100 μL of fresh sample (n=3) in 4.9 mL of 0.5 M hydrochloric acid for 1 hour before 50 μL was mixed with 2.45 mL of ferrozine solution (1 g ferrozine, 11.96 g (4-(2-hydroxyethyl)-1-piperazineethanesulfonic acid/L; adjusted to pH 7) in a cuvette (n = 3 per replicate). This was left to stand for exactly one minute before absorbance at 562 nm was measured using a spectrophotometer (Jenway 7315, Essex, UK). To quantify total bioavailable Fe (step 2), 200 μL of 6.25 M hydroxylamine hydrochloride was added to the digested samples and left to stand for another hour. This was then added to ferrozine solution in cuvettes and measured as before. Standards of known concentrations of FeSO_4_.7H_2_O were measured to allow conversion of absorbance to Fe concentrations.

### Copper growth assay

To confirm that metal concentrations in our river water and sediment samples were sufficiently high to select for metal resistance mechanisms, and to test whether liming impacted this selection, we used a copper growth assay. Specifically, we added 20 μL of either the ancestral *P. aeruginosa* strain or defrosted samples of the evolved populations to a 96-well plate well containing 180 μL of plain Iso-Sensitest broth (Oxoid) and 20 μL to a well containing 180 μL of Iso-Sensitest broth at a concentration of 1 g/L of copper sulphate (CuSO_4_; Alfa Aesar, Massachusetts, United States). The optical density OD_600_ was then read every 10 minutes for 18 hours using a Biotek Synergy 2 spectrophotometer. We used 1 g/L of copper sulphate as this equates to a copper concentration (6.26 mM) previously found in highly polluted environments (72, 73).

### Siderophore (CAS) assay

Total siderophore production was quantified using the Chrome Azurol S (CAS) assay (74). Samples were plated onto tryptic soy agar (TSA: Oxoid) supplemented with nystatin (Sigma: 20 μg/mL) to suppress fungal growth and X-gal. After 48 hours, *P. aeruginosa* colonies were counted to quantify density, before 24 colonies per replicate were randomly picked using sterile toothpicks. Selected colonies were resuspended in 1 mL of KB media in a deep 96-well plate and grown overnight at 28 °C. These were mixed with glycerol (final concentration 25%) and frozen at -80 °C. A scraping from each frozen monoculture was then grown in 2 mL of iron-limited casamino acid (CAA: Fisher) medium overnight. Iron limitation was caused by the addition of human apotransferrin (100 mg/mL; BBI Solutions, Crumlin) and sodium bicarbonate (20 mM; Acros Organics) to induce the production of siderophores. Cultures were centrifuged, and the supernatant assayed using the liquid CAS assay to quantify total siderophores, whilst pyoverdine was quantified by measuring the fluorescence of each culture at 460 nm following excitation at 400 nm. By measuring the optical density of the pre-centrifuged cultures and quantifying absorbance of sterile media, siderophore production per clone was estimated using: [1 − (Ai/Aref)] /[ODi)], where ODi = optical density 600 nm and Ai = absorbance at 630 nm of the assay mixture and Aref = absorbance at 630 nm of reference mixture (CAA+CAS) (18, 75).

### Galleria mellonella virulence assay

The insect infection model *Galleria mellonella* was used to quantify *P. aeruginosa* virulence (59, 60). Defrosted freezer stocks containing the whole sample microbiome were diluted 100-fold using M9 salt buffer, before 10 μL was injected into twenty final instar larvae per replicate using a 50 μL syringe (Hamilton, Nevada, USA). Injected larvae were incubated at 37 °C and mortality was monitored hourly after 13 hours for 12 hours with a final check at 42 hours. Larvae were classed as dead when mechanical stimulation of the head caused no response (60). M9-injected and non-injected controls were used to confirm mortality was not due to injection trauma or background *G. mellonella* mortality; >10% control death was the threshold for re-injecting (no occurrences). Prior to assays on microcosms containing *P. aeruginosa*, we confirmed the natural microbial community caused zero mortality by injecting replicates not inoculated with *P. aeruginosa* as described above.

### Antibiotic resistance assay

To test the evolved resistance of *P. aeruginosa* to the clinically relevant antibiotics apramycin, cefotaxime and trimethoprim, we used the same *P. aeruginosa* colonies isolated for the siderophore analysis. We first determined the minimum inhibitory concentration of the three antibiotics for our ancestral strain, by growing the ancestral strain for 24 hours (as described above), and plating it on TSA containing a range of concentrations of the antibiotics that increased in 10 μg/mL increments from 0 - 60 μg/mL. The minimum inhibitory concentrations were found to be 12 μg/mL, 30 μg/mL and 40 μg/mL for apramycin, cefotaxime and trimethoprim, respectively. Next, the individual evolved clones were defrosted before 2 μL of each was plated onto either plain TSA, TSA containing apramycin (Sigma: 15 μg/mL) cefotaxime (Molekula: 50 μg/mL) or TSA containing trimethoprim (Sigma: 60 μg/mL). Strains were considered resistant if colonies could be observed after 48 hours. The ancestral strain was used as a negative control.

### Statistical analysis

The effect of liming and shaking, plus their interaction, on the final pH, density of *P. aeruginosa* (log_10_(cfu mL^-1^)), and the proportion of total bioavailable iron (Fe^2+^ + Fe^3+^) that was Fe^2+^ (Fe^2+^ / (Fe^2+^ + Fe^3+^)) was tested using linear models with liming and shaking as explanatory variables. In general, model reduction was carried out by sequentially removing terms from the full model and comparing model fits using *F*-tests; we report parameter estimates of the most parsimonious model. The effect of pH on the density of *P. aeruginosa* populations was tested using a linear model with density (cfu mL^-1^) log_10_ transformed.

To test whether evolved samples had greater resistance to copper than the ancestral strain, we first calculated the relative fitness, *w*, of each population by dividing its maximum optical density after 18 hours when grown with copper (OD_maxC_) by its maximum optical density when grown without copper (OD_maxWC_), *i*.*e. w* = OD_maxWC_ / OD_maxC_. We then carried out a one way ANOVA with *w* as the response variable and treatment (including ancestor) as the explanatory variable. Secondly, we carried out a Dunnett’s test, using the ‘*DescTools’* R package (76), to test whether each treatment differed to the ancestor. Finally, we tested the effect of liming and shaking on the metal resistance of the final populations in a linear model, with log(*w*) as the response variable, and liming, shaking and their interaction as the explanatory variables. In all tests *w* was log transformed to normalise the residuals.

To test liming and shaking effects on total siderophore and pyoverdine production, linear mixed effects models (LMEM) were carried out using the ‘*lme4’* package (77) with liming and shaking as explanatory variables, and random intercepts fitted for each replicate to control for multiple clones being sampled from the same microcosm. For these LMEMs, we used the ‘*DHARMa’* package (78) to check residual behaviour, after which the most parsimonious model was arrived at by comparing models with and without the liming-shaking interaction using *χ*-tests. Two samples had pyoverdine values much lower than the rest, so a Grubbs test (*‘outliers’* package; (79)) was used to check if they were significant outliers. They were and therefore were removed from this and all further models to improve model fit. To test the association between copper resistance and both total siderophore and pyoverdine production, we carried out two linear models with log(*w*) as the dependent variable and either mean total siderophore production per microcosm or mean pyoverdine production per microcosm as the explanatory variable.

Virulence was analysed in three separate models. First, we tested whether larvae that died before 42 hours were injected with samples containing more siderophores and pyoverdine than those that remained alive after 42 hours. This was done by carrying out two separate binomial generalised linear mixed models (GLMM) using the ‘*lme4’* package (77), with number of *G. mellonella* dead versus alive as the binomial response variable, and either the production of total siderophores or pyoverdine as the explanatory variable. In this model pyoverdine production was log_10_ transformed to normalise residuals. Secondly, we tested whether the mean time it took deceased larvae (20 per replicate) to die was associated with total siderophore and pyoverdine production (both values taken from the mean of 24 clones) using a linear model. Finally, we tested whether virulence differed between treatments. To do this, survival curves were fitted using Bayesian regression in the R package ‘*rstanarm*’ (80) and the package ‘*tidybayes*’ (81) was used to estimate parameters. A proportional hazards model with an M-splines baseline hazard was fitted, with liming, shaking plus their interaction as fixed effects. We additionally included random intercepts for each sample to control for multiple (20) *G. mellonella* being inoculated with the same sample. Models used three chains with uninformative priors and were run for 3000 iterations. Model convergence was assessed using Rhat values (all values were 1), before we manually checked chain mixing.

The proportion of apramycin, cefotaxime, and trimethoprim resistance in each treatment (number of resistant colonies out of 24 in total) was compared using Kruskal-Wallace non-parametric tests, with resistance proportion as the response variable and treatment as the explanatory variable. All analyses were carried out in R version 3.3.3 (82).

## Results and discussion

### Liming and shaking affected pH but not the relative abundance of Fe^2+^

Here, we tested whether liming of metal-contaminated aquatic environments decreases co-selection for virulence and antibiotic resistance in the opportunistic pathogen *P. aeruginosa*. To do this, we evolved *P. aeruginosa* with or without lime in microcosms containing a mixture of metal contaminated river water and sediment in the presence of the natural microbial community. We employed both shaking and static microcosms to represent turbulent and stagnant aquatic environments, in order to test whether liming effects were dependent on environmental structure (Fig. 1).

As expected, liming significantly decreased the acidity of sediment and water from the initial pH of 5.8. However, the extent of this effect was significantly greater in the shaking treatments (liming-shaking interaction: F_1,20_=23.1, p<0.001; Fig. 2), likely due to increased mixing of lime and oxygen throughout the microcosms. The shaken-limed treatment reached a pH of 7.2 (+/- 0.11 SD) whereas the static-limed treatment reached a pH of 6.7 (+/- 0.25 SD). Both non-limed treatments had a final pH of 5.7 (+/- 0.19 SD). As pH is often a good predictor of iron speciation (83), we tested how the treatments affected the relative proportions of Fe^2+^ and Fe^3+^. We found the proportion of more bioavailable Fe^2+^ to not significantly differ as a result of liming, shaking, nor their interaction (lime main effect: F_1,9_=3.47, p=0.10; shaking main effect: F_1,9_=3.00, p=0.12; lime-shaking interaction F_1,8_=0.73, p=0.42; Fig.2), with Fe^2+^ making up 82% of the total iron available on average across the treatments. Given that iron speciation remained similar in all treatments, this indicates that the redox potential within the microcosms did not change to become more anaerobic under static conditions (83).

**Figure 2.**
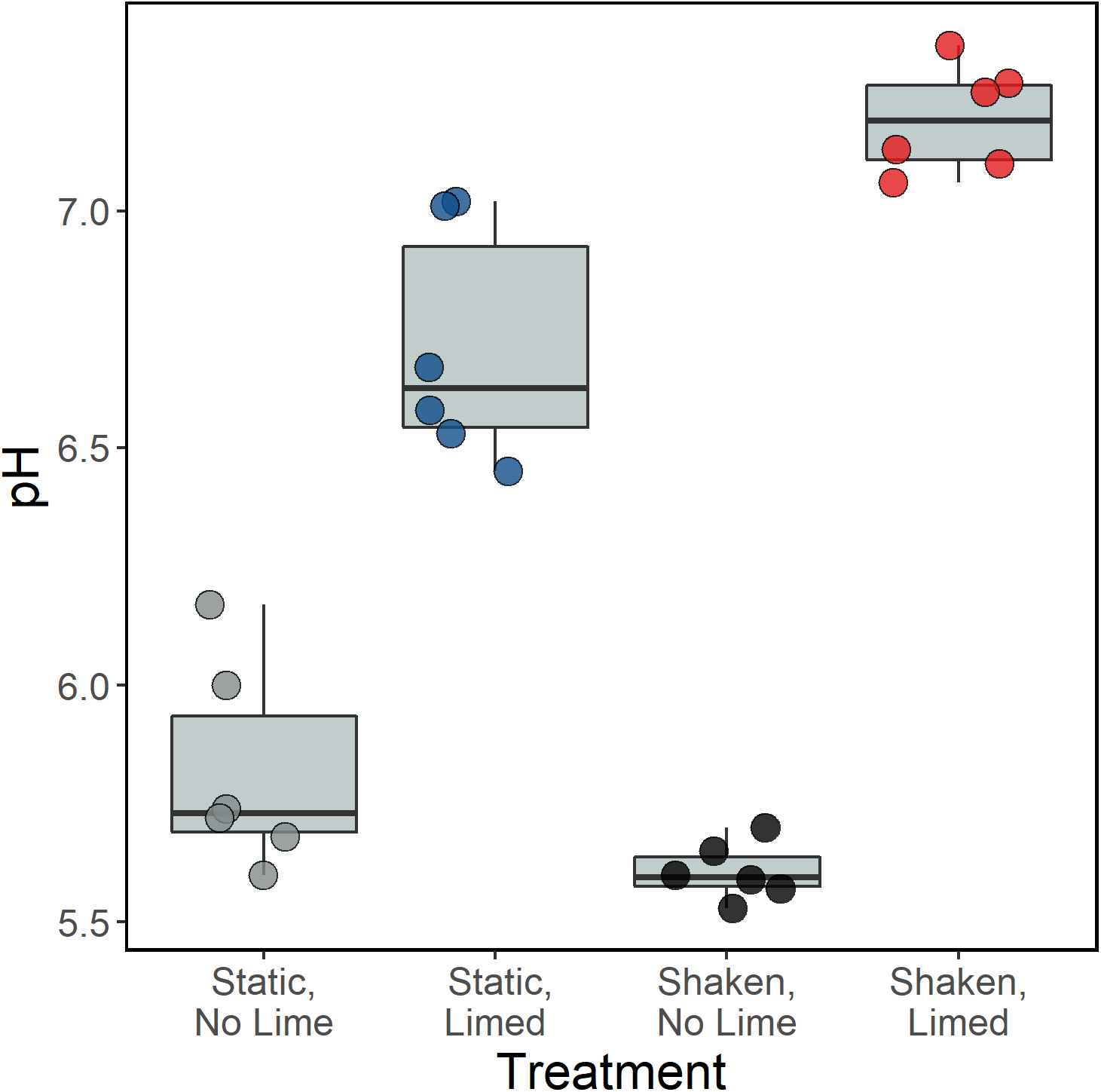
The final pH of microcosms containing river water and sediment after 28 days incubation. We used a factorial design with limed and shaken treatments, each with six replicates (each represented by a white circle). The starting pH was 5.8. The significant effect of liming on pH (p<0.001) was increased through an interaction with shaking (p<0.001).

Hence iron bioavailability was not significantly influenced by the different experimental conditions and therefore iron limitation was unlikely to represent a significant driver for siderophore production.

### P. aeruginosa populations incubated without lime had greater tolerance to copper

In order to test whether our river water and sediment samples selected for greater metal resistance, we incubated the ancestral *P. aeruginosa* strain and final populations in a medium containing a high concentration of copper (1 g/mL of copper sulphate). We then compared the maximum optical density of each culture relative to that of cultures grown without copper (*w*). Confirming that our samples contained toxic metals, the ancestral strain had a lower relative fitness (*w*) when grown with copper than all final populations (Dunnett’s test: p=<0.013 for all contrasts; Fig. 3). Moreover, when comparing the effect of the different treatments on *w*, we found populations from the non-limed treatments to have greater relative fitness in a toxic copper environment than those from the limed treatments (liming main effect: F_1,21_=4.44, p=0.047; Fig. 3), but shaking to not have an effect on relative fitness (shaking main effect: F_1,21_=0.83, p=0.074; liming-shaking interaction: F_1,20_=0.11, p=0.75). These results demonstrate that our river microcosms imposed strong selection on *P. aeruginosa* to evolve metal resistance, and that liming reduced this selection pressure.

**Figure 3.**
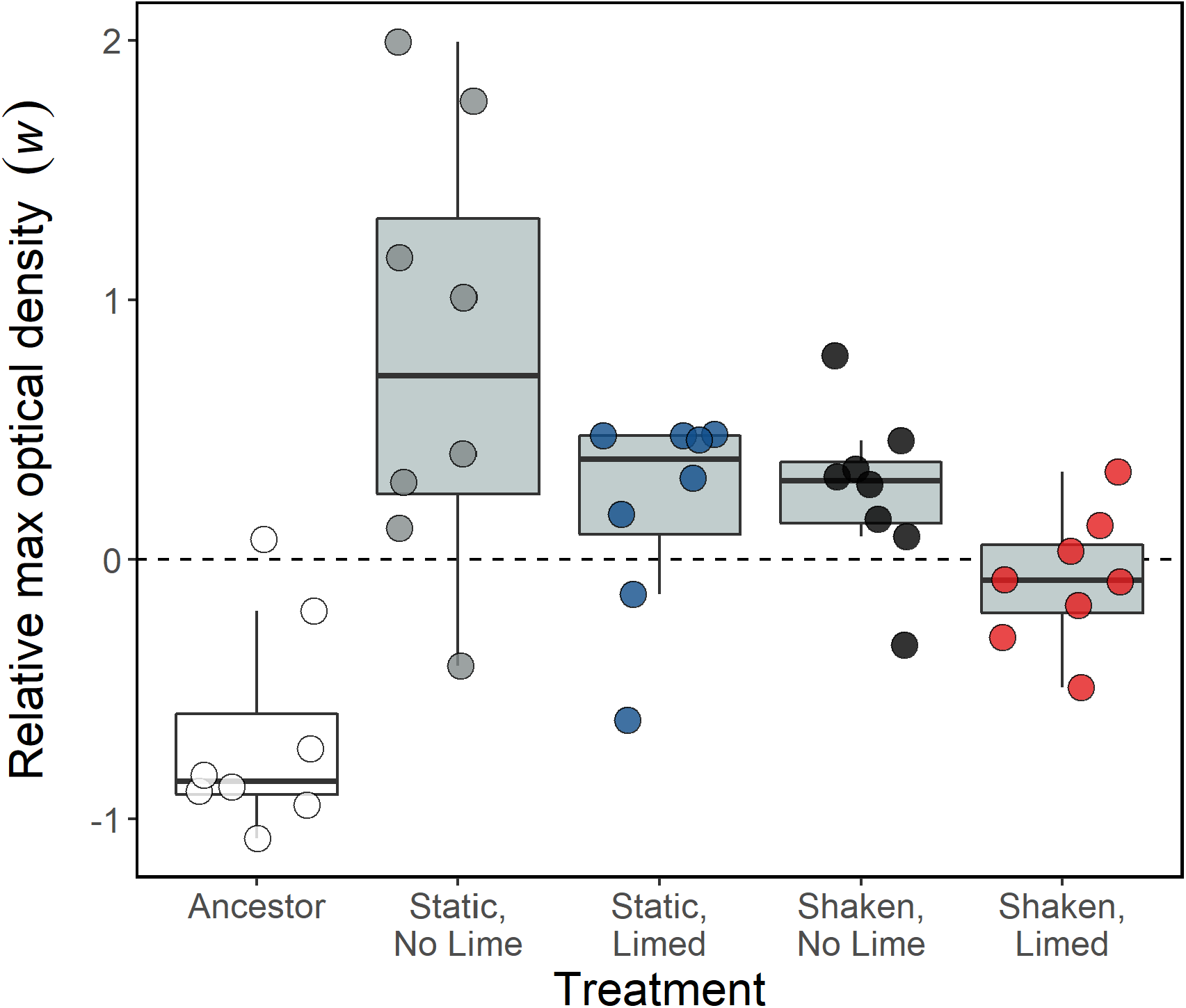
The maximum optical density (OD_600_) of *P. aeruginosa* populations after 18 hours incubation with toxic copper (1g/L copper sulphate) relative to their maximum OD_600_ when grown without copper (log *w*). Populations above the horizontal dashed line at 0 (log of 1) have a higher relative max OD_600_ when grown with copper, whereas those below the line have reduced maximum growth. The white box shows the ancestral strain, whereas the grey boxes show populations incubated in microcosms containing river water and sediment for 14 days. Circles show individual replicates (n=6), and colours the different treatments.

### Neither liming nor shaking affected P. aeruginosa density or siderophore production

Next, we tested the treatment effects on *P. aeruginosa* density and siderophore production. The final density of *P. aeruginosa* after two weeks of evolution varied substantially between samples (1.1 × 10^6^ ± 1.6 × 10^6^ SD cfu/mL), but was not significantly affected by liming, shaking, nor their interaction (liming main effect: F_1,21_=1.96, p=0.18; shaking main effect: F_1,21_=2.77, p=0.11; liming-shaking interaction: F_1,20_=0.70, p=0.41). There was also no significant effect of pH on *P. aeruginosa* density (F_1,22_=0.97, p=0.36). Although pH can affect bacterial density (84), our finding of no effect is consistent with previous results demonstrating that *P. aeruginosa* densities are similar across an equivalent pH range as used here (85).

To test whether liming and shaking affected siderophore production, both total siderophore production and the production of pyoverdine — the primary siderophore produced by *P. aeruginosa* (56) — were measured for 24 clones per replicate (24 × 24 clones). Quantifying pyoverdine production in addition to total siderophores is important, as it is a key virulence factor in *P. aeruginosa* but its production does not necessarily correlate with that of other siderophores, such as pyochelin (62). We found neither liming, shaking nor their interaction significantly affected mean total siderophore production (liming main effect: *χ*^*2*^=1.49, d.f.=1, p=0.22; shaking main effect: *χ*^*2*^=0.49, d.f.=1, p=0.48; liming-shaking interaction: *χ*^*2*^=0.08, d.f.=1, p=0.78) or pyoverdine production (liming main effect: *χ*^*2*^=0.56, d.f.=1, p=0.46; shaking main effect: *χ*^*2*^=2.3, d.f.=1, p=0.13; liming-shaking interaction: *χ2*=1.14, d.f.=1, p=0.29). However, we note that there was a large variation in production between the 24 clones used to represent each microcosm (mean production: total siderophores = 4.23; pyoverdine = 766; *σ*_replicate_: total siderophores = 1.94; pyoverdine = 69.3), and that two pyoverdine values were significant outliers and consequently were removed from all further analysis in order for model assumptions to be met (these were one from the non-limed shaken treatment (pyoverdine production = 26.9, p<0.001) and one from the limed-static treatment (pyoverdine production = 174, p<0.001); which were lower than the pre-removed mean pyoverdine production of 710.6 and median of 789.8).

That siderophore production, which is regulated by iron availability and the presence of toxic metals, did not significantly differ between treatments concurs with the non-significant differences in Fe^2+^ availability between treatments. However, it is surprising that siderophore production was not reduced by liming, given that *P. aeruginosa* populations from the limed treatments were less tolerant to toxic copper. To explore this further, we tested whether either total siderophore or pyoverdine production was associated with copper tolerance, and found neither of them to be (Total siderophores: F_1,20_=0.013, p=0.91; Pyoverdine: F_1,20_=0.294, p=0.59). This suggests other metal resistance mechanisms, such as decreased outer membrane permeability and increased induction of ATPase efflux transporters, could be responsible for the increased copper tolerance of evolved populations (86). Our finding of no significant differences in siderophore production contrasts with that of Hesse and co-workers (18), who found that the addition of lime to soils collected in the near vicinity of our locality significantly reduced community-wide siderophore production. This difference is most likely due to shifts in siderophore production driven by changes in community composition with liming selecting for non-producing isolates (18), whereas here we solely focused on siderophore production by *P. aeruginosa*. This suggests that although liming reduces community-wide siderophore production in metal-contaminated acidic soils, this effect may not be seen in specific species. Interestingly, *P. aeruginosa* has been proposed as a suitable siderophore-producing bacterium for use in phytoremediation, which relies on the combined use of microorganisms and plants to aid toxic metal remediation (87, 88). It has been proposed that liming, by reducing siderophore production, may hinder phytoremediation (18) as metal-uptake by plants is often increased when metals are bound to bacterial siderophores. Given that no significant effect of liming on siderophore production by *P. aeruginosa*, was observed, we suggest that liming and *P. aeruginosa-*assisted phytoremediation could be used simultaneously without compromise.

### Virulence did not differ between treatments, but was positively associated with siderophore production

As we found a large variation in siderophore production, which is a known virulence factor in *P. aeruginosa* (64), we tested whether virulence, quantified using the *G. mellonella* infection assay, differed as a consequence of pyoverdine production, total siderophore production or treatment. Firstly, we tested whether *G. mellonella* larvae alive at the final time check (42 hours) had been injected with populations producing less total siderophores and pyoverdine compared with larvae that died before this point, and found that they were (total siderophores: *χ*^2^=6.11, d.f.=1, p=0.013; pyoverdine: *χ*^2^=6.98, d.f.=1, p=0.004). Next, we tested whether increased siderophore and pyoverdine production resulted in increased virulence. We found a significant positive association between virulence (mean time to death per population) and both total siderophore and pyoverdine production (total siderophores: F_1,22_=8.9, p=0.007; Fig. 4A; pyoverdine: F_1,20_=10.3, p=0.004; Fig. 4B), with every one unit increase in total siderophore production decreasing mean time to death by 1.42 hours (SE = 0.4745), and every one unit increase in pyoverdine production decreasing the mean time to death by 0.032 hours (SE = 0.0098). This positive association between siderophore production and virulence is supportive of previous work both in *G. mellonella* (37) and murine models (89). In addition to directly aiding iron uptake within a host, siderophores can increase virulence by triggering other virulence factors (90, 91). Finally, virulence was compared between treatments using survival curves (Fig. 4C). Virulence did not significantly differ as a function of treatment, with the credible intervals for liming, shaking and their interaction all crossing 1. No significant treatment effect on virulence is concurrent with the finding that the treatments did not significantly affect siderophore production. Finding virulence to not be significantly different between structured (static) and unstructured (shaking) environments contrasts with findings by Granato and co-workers (92), who found that pyoverdine-mediated virulence in *P. aeruginosa* was greater when grown in solid media than in liquid. The lack of changes detected in siderophore production and virulence between the experimental treatments might be due to the more subtle (and arguably more realistic) conditions under which spatial structure was varied in our study, as well as the presence of a resident microbial community.

**Figure 4.**
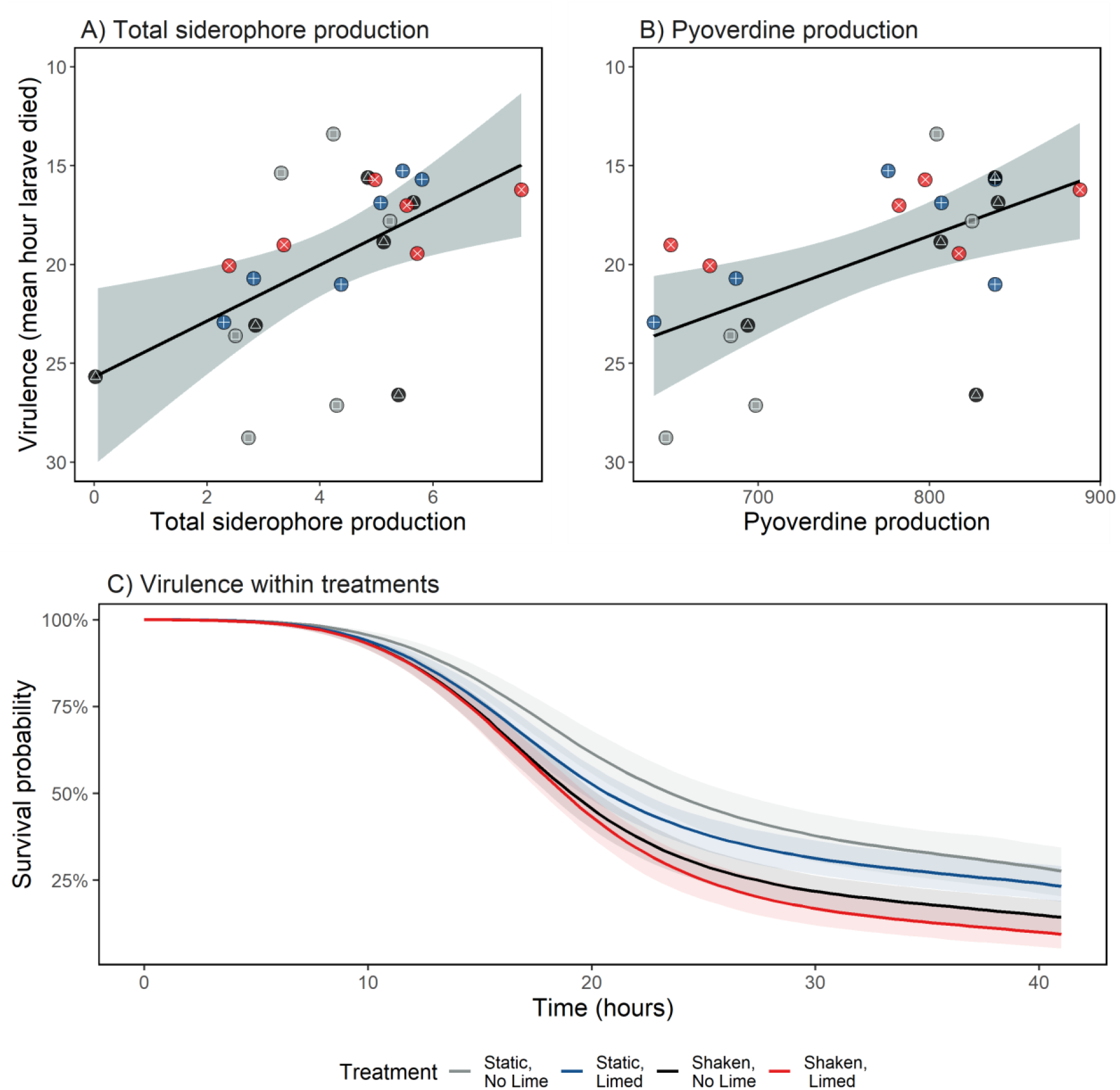
Mean virulence of *Pseudomonas aeruginosa* evolved in metal-contaminated aquatic communities as a function of (A) mean total siderophore production and (B) mean pyoverdine production. Virulence was quantified using the *Galleria mellonella* infection model (n = 20 per replicate) and given as the mean time to death. Pyoverdine and total siderophore production were measured in standardised fluorescence units per OD_600_. Individual circles show the mean production by 24 clones from each replicate. Colours and shapes represent different treatments: grey and □ = static, no lime, blue and + = static, limed, black and △ = shaken, no lime, and red and × = shaken, limed. Panel C shows the change in survival probability of larvae over time within each treatment. These do not significantly differ from one another. Shaded areas represent 95% confidence intervals.

### Antibiotic resistance evolution

As metal pollution has been shown to co-select for antimicrobial resistance (38), we tested whether lime addition altered *P. aeruginosa* resistance to the clinically relevant antibiotics apramycin (15 μg/mL), cefotaxime (50 μg/mL) and trimethoprim (60 μg/mL) after evolution in metal-contaminated river sediments. Increased resistance was observed in all treatments (Fig. 5), with neither lime nor shaking affecting resistance to any of the antibiotics tested (apramycin: chi-squared=2.35 p=0.50 df=3; cefotaxime: chi-squared=2.98 p=0.40 df=3; trimethoprim: chi-squared=5.25 p=0.16 df=3; Fig. 5). Of note, one sample from the shaken, non-limed treatment consistently had the lowest resistance to all three antibiotics, with no isolates from this population being resistant to cefotaxime or trimethoprim, and fewer than 50% being resistant to apramycin. Our observation of rapid evolution of antibiotic resistance in the other replicates and treatments supports existing evidence that metal contamination can pose an important co-selective pressure for resistance (44, 49, 93), including in *P. aeruginosa* (42). That resistance did not differ significantly between treatments in our experiment demonstrates that liming to pH ∼7 is not effective at remediating this co-selective effect, and neither was the loss of spatial structure via shaking. A plausible reason for this is that liming reduces metal bioavailability by precipitating ions from solution into the solid phase. This would mean cells in the sediment (the vast majority of the population) would still be exposed to metals where, although at a lower bioavailability, they can still be a cause of co-selection (48). Although we did not determine the mechanistic basis of co-selection, we note that cross-resistance, co-resistance and co-regulation mechanisms have all been reported for Pseudomonads, and the altering of cellular targets is a mechanism commonly used by *P. aeruginosa* to tolerate metal, trimethoprim and beta-lactam antibiotics such as cefotaxime (42, 94). We are aware of a single study testing the effects of liming on antimicrobial resistance (23). This study found liming decreased the susceptibility of Rhizobium species from soil to multiple antibiotics, and hypothesised this was due to a greater production of natural antibiotics at near-neutral pH selecting for resistance. Although we note that increasing soil pH will generally decreases the bioavailability of any metals present, the authors (23) stated that metal effects would not be operative in their study, suggesting no metal contamination was present.

**Figure 5.**
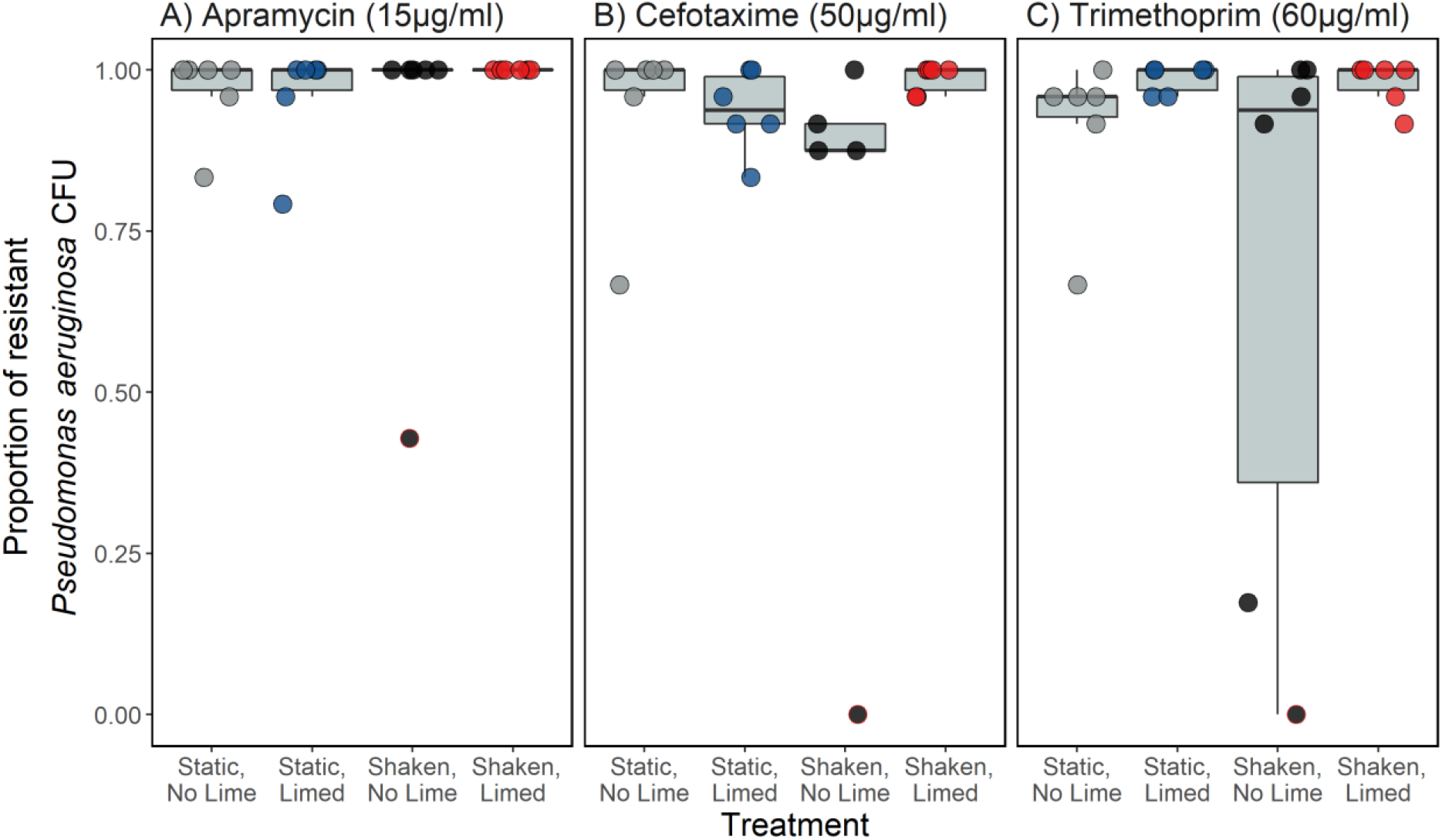
The proportion of 24 *Pseudomonas aeruginosa* clones per replicate (n=6) resistant to (A) apramycin (15 μg/mL), (B) cefotaxime (50 μg/mL) and (C) trimethoprim (60 μg/mL) antibiotics. Clones were tested after two weeks of evolution in microcosms containing metal-contaminated river water and sediment while embedded in the resident microbial community. Circles show individual replicates; those with a red outline are from the same sample, which is the least resistant to all three antibiotics.

## Conclusion

*P. aeruginosa* populations evolved metal resistance after two weeks, and liming reduced this effect. However liming and spatial structure (shaking) were observed to have little effect on *P. aeruginosa* pathogenic traits. Despite finding a positive association between siderophore production and virulence, neither siderophore production nor virulence systematically differed between treatments, suggesting that liming does not alter the effect of metals on siderophore-mediated virulence in *P. aeruginosa*. This finding also implies that concurrent use of liming and *P. aeruginosa-* assisted phytoremediation techniques is possible in scenarios were this bacterium can persist in a natural community. Moreover, we found *P. aeruginosa* rapidly evolved resistance to three clinically relevant antibiotics regardless of treatment. We therefore show that a common metal remediation method did not reduce metal pollution-based co-selection for virulence or antibiotic resistance. Importantly, these findings further our understanding of how key determinants of pathogenicity evolve outside of the clinical setting, and further demonstrate that metal-polluted environments can select for them.

